# Human motor unit discharge patterns reveal differences in neuromodulatory and inhibitory drive to motoneurons across contraction levels

**DOI:** 10.1101/2023.10.16.562612

**Authors:** Jakob Škarabot, James A Beauchamp, Gregory EP Pearcey

**Author notes:** All authors contributed to the work equally.

## Abstract

All motor commands converge onto motor units (MUs), which transduce the signals into mechanical actions of muscle fibres. This process is highly non-linear due to combinations of ionotropic (excitatory/inhibitory) and metabotropic (neuromodulatory) inputs. Neuromodulatory inputs facilitate dendritic persistent inward currents, which introduce non-linearities in MU discharge patterns and provide insights into the structure of motor commands. Here, we investigated the relative contribution of neuromodulation and the pattern of inhibition to modulate human MU discharge patterns with contraction forces up to 70% maximum. Leveraging MU discharge patterns identified from three human muscles (tibialis anterior – TA, and vastus lateralis and medialis), we show that with increased contraction force, the onset-offset discharge rate hysteresis (ΔF) increased whilst ascending MU discharge patterns become more linear, with lower slopes. In a follow-up experiment, we demonstrated that the observations of increased ΔF and more linear ascending MU discharge patterns with greater contraction force are maintained even when accounting for contraction duration and rate of force increase. We then reverse-engineered TA MU discharge patterns using highly realistic in silico motoneuron pools to substantiate the inferred physiological mechanisms from human recordings. We demonstrate a sharply restricted solution space, whereby the contraction force-induced changes in experimentally obtained MU discharge patterns can only be recreated with increased neuromodulation and a more reciprocal (i.e. push-pull) inhibitory pattern. In summary, our experimental and computational data suggest that neuromodulation and inhibitory patterns are uniquely shaped to generate discharge patterns that support force increases across a large proportion of the motor pool’s recruitment range.

**Significance statement:** How the structure of motor commands is modified to scale motor output is largely speculative despite its critical role in the neural control of movement. Here, we demonstrate that human motor unit discharge patterns become more linear and exhibit greater discharge rate hysteresis with greater contraction force. These experimentally observed patterns can only be replicated in silico with biophysical models of spinal motoneurons by increasing neuromodulation and shifting inhibitory commands to be more reciprocal to excitation (i.e., push-pull excitation-inhibition synaptic control). Collectively, these results suggest that the structure of motor commands is uniquely orchestrated to support increases in contraction force.

## INTRODUCTION

The transformation of motor commands into mechanical actions of muscle fibres via the motor unit (MU, alpha motoneuron and its innervated muscle fibres) is a non-linear function influenced by both ionotropic and metabotropic inputs. Ionotropic inputs (e.g., corticospinal, reticulospinal, sensory, etc.) provide excitation (depolarisation) and inhibition (hyperpolarisation) to motoneurons, whilst metabotropic (neuromodulatory) inputs modulate ionotropic inputs via G-protein coupled receptors. One potent source of neuromodulation is monoamines (serotonin (5-HT) and noradrenaline), which are released via long descending axons from the raphe nuclei and locus coeruleus that project diffusely throughout the spinal cord (Goaillard and Marder, 2021), and facilitate motoneuronal persistent inward currents (PICs) mediated by voltage-gated ion channels (Schwindt and Crill, 1980; Hounsgaard et al., 1988; Bennett et al., 1998). These PICs provide an additional source of depolarisation that introduces non-linearities in motoneuron discharge by amplifying and prolonging excitatory ionotropic inputs (Lee and Heckman, 1998a, 2000).

Whilst the importance of modulating excitatory ionotropic inputs for encoding muscle contraction force has received significant attention in both primates (Cheney and Fetz, 1980; Glover and Baker, 2022) and humans (Oya et al., 2008; Weavil et al., 2015), the contribution of neuromodulatory inputs to greater force production is less clear. Evidence in the cat indicates that 5-HT release is tightly coupled with the intensity of motor output (Jacobs et al., 2002) and pharmacological studies in humans suggest the gain-control for voluntary force production is mediated by 5-HT (Wei et al., 2014; Kavanagh et al., 2019). Nevertheless, best estimates of neuromodulatory inputs in humans (onset-offset discharge hysteresis of MU pairs, ΔF; Gorassini et al., 2002a) show inconsistent results, with ΔF either increasing (Orssatto et al., 2021; Goodlich et al., 2023) or remaining constant (Afsharipour et al., 2020; Kim et al., 2020) with increased voluntary drive. Notably, however, prior studies examining human motoneuron properties inferred from MU discharge patterns have been largely restricted to relatively low contraction forces (≤30% of maximum), which limits the investigation to predominantly lower threshold MUs and restricts the range of behaviour to conditions where intense neuromodulation may not be required to facilitate force output.

Though PICs are proportional to monoamines, they are also highly sensitive to local inhibitory inputs; that is – tonic inhibitory inputs (Hyngstrom et al., 2007) as well as the pattern of inhibition relative to excitation (e.g., reciprocal/push-pull vs. proportional patterns; Johnson et al., 2012, 2017a). Tonic inhibitory input linearises human MU discharge patterns (Revill and Fuglevand, 2017) and reduces estimates of PIC magnitude in humans (Mesquita et al., 2022; Pearcey et al., 2022), in agreement with prior studies in cat (Kuo et al., 2003). Moreover, realistic simulations of motoneuron pools suggest that ΔF is likely to be greater with reciprocal (i.e., push-pull) rather than proportional inhibitory pattern (Beauchamp et al., 2023; Chardon et al., 2023), an effect that has largely been neglected in prior studies investigating the influence of contraction level on MU discharge rate profiles.

Here, we performed two experiments in humans, followed by computer simulations, to characterise human MU discharge patterns across a wide range of contraction levels. In Experiment 1, we characterised MU discharge rate pattern modifications across a range of contraction levels in three different human muscles (tibialis anterior, TA; vastus lateralis and medialis, VL and VM). In Experiment 2, we investigated how the duration and rate of the force increase influenced MU discharge rate pattern modifications across force levels. We then used the experimentally obtained data to recreate MU discharge patterns using highly realistic motoneuron models to elucidate the relative contribution of neuromodulation and inhibitory input patterns to the MU discharge rate modifications across force levels. We hypothesised that 1) estimates of PICs would increase with contraction level; 2) MU discharge rate patterns would indicate a more reciprocal pattern of inhibition relative to excitation (i.e., push-pull) with greater contraction level; and 3) the contraction- level dependent effects would be maintained irrespective of the rate of force increases and duration of contractions.

## MATERIALS AND METHODS

### Human experiments

#### Participants

Two human experiments were performed; 15 healthy adults participated in Experiment 1 (4 females; age: 24 ± 5 years, stature: 1.77 ± 0.08 m, mass: 71.6 ± 12.3 kg) and 11 healthy adults participated in Experiment 2 (3 females; age: 25 ± 4 years, stature: 1.77 ± 0.08 m, mass: 78.2 ± 13.8 kg). Participants had no known cardiovascular, neuromuscular, or musculoskeletal impairments, and did not take medications known to affect the nervous system. The study was approved by Loughborough University ethics committee (2021-5361-4724) in accordance with the latest version of Declaration of Helsinki, except for registration in database. Participants provided written, informed consent before taking part in any experimental procedures.

### Human experimental design

#### Experiment 1

Participants visited the laboratory on three occasions, separated by at least 48 hours and no longer than 10 days. The first visit involved familiarisation with the performance of submaximal and maximal dorsiflexion and knee extension. Participants then attended two experimental sessions during which they performed isometric knee extension and dorsiflexion, respectively (randomised order). Each experimental session started with a series of warm-up submaximal isometric contractions (3 x 50, 3 x 75, and 1 x 90% of perceived maximal voluntary force; MVF). Participants then performed two to three contractions with maximal effort, separated by one minute rest, and the best trial was considered their MVF. Following the determination of MVF, participants performed triangular contractions at several force levels: 15, 30, 50, and 70% of MVF. Two contractions were performed at each level. The order of contraction levels was pseudorandomised; for contractions at 15- 50% MVF, the order was randomised, with the 70% MVF contractions always performed last to minimise the decline in contractile function. For each contraction, participants linearly increased and decreased force to/from the target (i.e., peak) force level across 10 s in each direction (i.e., 20 s total). The contraction duration was kept constant (i.e., 20 s total) across all contraction levels, resulting in rate of force increase/decrease of 1.5, 3, 5 and 7% MVF/s respectively. To minimise MU recruitment threshold accommodation (i.e., warm-up; 35), at least 30, 45 and 90 seconds of rest was provided between contractions performed at 15-30, 50, and 70% MVF, respectively.

#### Experiment 2

Both contraction duration and rate of force increase can alter recruitment-derecruitment hysteresis through spike-frequency adaptation and spike-threshold accommodation, respectively, as indicated by simulations (Revill and Fuglevand, 2011). Spike-frequency adaptation, which denotes a time-dependent decrease in MU discharge via inactivation of sodium channels and calcium-dependent potassium conductance (Sawczuk et al., 1995; Powers et al., 1999; Miles et al., 2005a), appears to be particularly influential in contaminating the ΔF estimation, with a greater influence compared to spike-threshold accommodation (Vandenberk and Kalmar, 2014). Thus, we prioritised standardisation of contraction duration over the rate of force increase in Experiment 1. Nevertheless, spike-frequency adaptation and spike-threshold accommodation share some of the same mechanisms; for example, sodium channel inactivation has been shown to contribute to both phenomena in rat preparations (Bradley and Somjen, 1961; Miles et al., 2005b). To assess the potential influence of spike-threshold accommodation and spike-frequency adaptation on our results, we performed an additional experiment in the dorsiflexors. Here, participants were tasked to perform contractions of several submaximal intensities (30, 50, 70% MVF), but they did so by either performing contractions of the same duration (10 s increase/decrease, 20 s in total; rates of force increase/decrease: 3, 5, 7% MVF/s) or the same rate of force increase/decrease (5% MVF/s; total contraction durations: 12, 20, 28 seconds). Two contractions were performed per each condition and the same rest periods were employed between contractions as in Experiment 1.

### Human experimental procedures

#### Force signals

To record dorsiflexion forces, participants were seated on a custom-made chair fitted with an ankle ergometer (NEG1, OT Bioelettronica, Torino, Italy) connected to a strain gauge (CCT Transducer s.a.s., Torino, Italy), the signal of which was amplified (×200; Forza-B, OT Bioelettronica, Torino, Italy). The ankle of the dominant leg was positioned at 10° of plantar flexion (0° = anatomical position), the hip was flexed at 120° (180° = full extension) and the knee was fully extended. The foot was strapped to the dynamometer with Velcro at tarsometatarsal and metatarsal phalangeal joints. The leg was additionally strapped over the knee joint to minimise its flexion. Knee extensor forces were recorded with participants seated on a custom-made isometric dynamometer with the knee and hip flexed at 115 and 125° (180° = full extension). The dominant leg was strapped (35-mm wide canvas webbing) at 15% of the tibial length in series with the strain gauge (amplification: ×370; Force Logic, Swallowfield, UK). The force signals for both dorsiflexion and knee extension were sampled at 2048 Hz and digitised with a multichannel amplifier (16-bit, Quattrocento; OT Bioelettronica, Torino, Italy).

#### High-density surface electromyography

High-density surface electromyograms (EMG) were recorded by placing a grid of 64 electrodes (13 rows x 5 columns, with one missing electrode; 1 mm electrode diameter, 8 mm inter-electrode distance; GR08MM1305, OT Bioelettronica, Torino, Italy) via disposable bi-adhesive foam layer (Spes Medica, Battipaglia, Italy) with its cavities filled with conductive paste (Ten20, Weaver and Company, Aurora, CO, USA) over the muscle bellies of tibialis anterior, vastus lateralis, and vastus medialis, with longitudinal axis of the electrode grid aligned with the presumed orientation of muscle fibres. The reference electrode (3M Red Dot, 3M Deutschland GmbH, Germany) was placed on the medial malleolus and the patella when recording the signals during dorsiflexion and knee extension, respectively. A dampened strap electrode was placed around the ankle of the non-dominant leg to ground the signal. The high-density EMG signals were recorded in monopolar configuration, band-pass filtered (10-500 Hz), digitised (2048 Hz) using a multichannel amplifier (16-bit, Quattrocento; OT Bioelettronica, Torino, Italy), and acquired using OT Biolab+ software (OT Bioelettronica, Torino, Italy).

### Human experimental data analysis

#### Force signal

In the process of off-line analysis, a 20 Hz low pass filter (4^th^ order, zero-lag, Butterworth) was applied to the force signal to remove non-physiological properties of the signal. The force offset was removed (i.e., gravity correction) before further analysis.

### High-density surface electromyography

#### Decomposition

Monopolar high-density surface EMG signals were initially inspected, and the channels deemed of poorer quality based on area under power spectrum and amplitude were removed (typically 2-3 channels). Monopolar signals were decomposed into individual MU spike trains using the Convolution Kernel Compensation algorithm (Holobar and Zazula, 2007). This extensively validated algorithm inverts the EMG mixing model and estimates the separation vectors (MU filters) that represent weights of the linear spatial-temporal EMG channel combinations that yield the estimation of individual MU spike trains (Holobar et al., 2010, 2014). Signals from each contraction were decomposed independently. Decomposition results were visually inspected by an experienced operator who iteratively optimised MU filters using procedures described previously (Del Vecchio et al., 2020). Only the results with regular discharge patterns and/or those exhibiting the pulse-to-noise ratio ≥ 30 dB (Holobar et al., 2014) were retained (accuracy >90%, false alarm rate <5%).

### Motor unit tracking

We examined MU discharge patterns with and without consideration for potential MU replicates across contraction levels (see below). For the former, it was important to establish which MUs were identified across the examined contraction levels. To identify unique MUs, we used the separation vectors (MU filters), to identify MU spike trains that were common across contraction levels. The reader is referred to previously published work on the mathematical basis and validation of this approach (Francic and Holobar, 2021; Škarabot et al., 2023). Briefly, the decomposition results of individual contractions performed at different intensities were concatenated. Considering the difficulty in identifying lower threshold MUs in the presence of much higher threshold MUs, we only concatenated contractions performed at 30, 50, and 70% MVF. In other words, we opted to exclude the trials performed at 15% MVF in this procedure as the likelihood of identifying a MU recruited during both 15 and 70%

MVF was very low (Francic and Holobar, 2021). For each contraction level and individual, we selected the contraction trial with the greatest MU yield after decomposition and/or the best fit to the triangular target force. The estimated MU filters from signals obtained during one contraction were then transferred to other contractions in the concatenated signal to identify common MUs (Francic and Holobar, 2021). Since duplicate MU spike trains can be identified when transferring MU filters, we identified those sharing 30% of the same MU discharges (discharge tolerance of 0.5 ms), and then removed them by discarding the duplicate with the lower estimated decomposition accuracy (i.e., lower pulse-to-noise ratio). As shown previously, the success of MU filter transfer varies across contraction intensities, with greater difficulty identifying discharges of lower-threshold MUs in the presence of higher-threshold MUs (Francic and Holobar, 2021). To maximise the number of tracked MUs, we therefore tracked MUs either across all concatenated contraction intensities (30-50-70% MVF) or only between neighbouring contractions (30-50 or 50-70% MVF; Figure 1).

**Figure 1.**
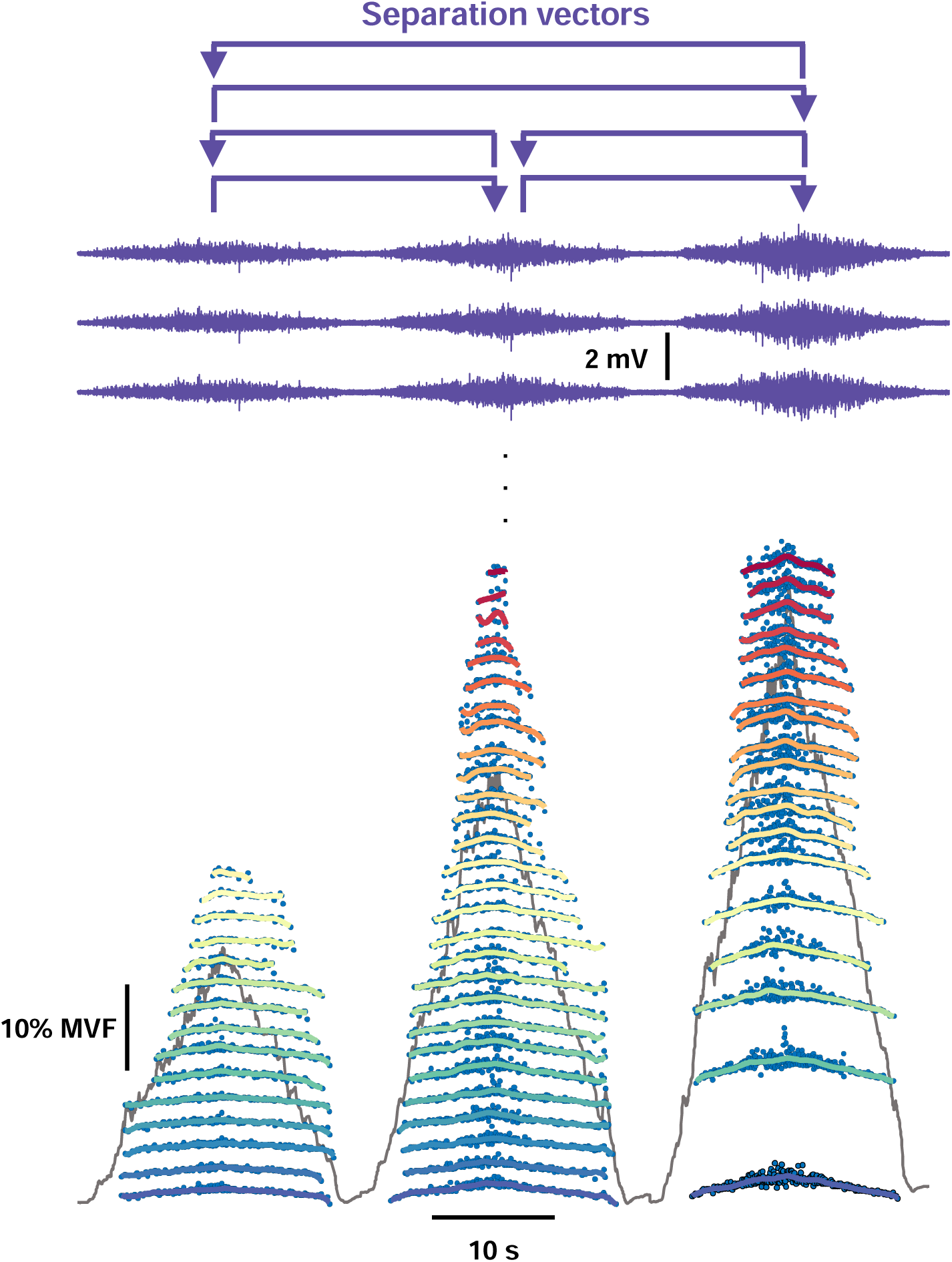
Identification of motor unit discharges. The approach to tracking motor units (MUs) across contraction intensities involved the estimation and application of separation vectors (MU filters) among all the possible combinations from signals obtained during contractions to 30, 50 and 70% of maximal voluntary force (MVF). Discharge patterns of tracked MUs smoothed with support vector regression are shown in colours. For clarity, only three channels of monopolar electromyography (EMG) signals are displayed.

### Separation vectors

#### Motor unit discharge rate metrics

##### Processing of motor unit spike trains

Support vector regression (SVR) was used to generate continuous estimates of MU discharge behaviour from the binary MU spike trains using procedures described previously (Beauchamp et al., 2022). Briefly, the instantaneous discharge rate for each MU was calculated using the reciprocal of its interspike interval, which was used to train an SVR model and generate the smooth estimates of discharge rate between recruitment and derecruitment using previously suggested parameters (Beauchamp et al., 2022).

##### Recruitment threshold and discharge rate

Recruitment and derecruitment thresholds were estimated as the instantaneous force at the instant of the first and last spike in the binary spike train, respectively. Peak discharge rate was calculated as the maximal value of the smoothed estimate of MU discharge, whereas onset and offset discharge rate were calculated as the first and last point of the SVR smoothed discharge.

##### Quantification of discharge rate hysteresis with respect to peak force

To quantify the discharge rate hysteresis on a single MU level, we calculated the ascending and descending duration of MU discharge with respect to peak force, and then calculated the difference in the duration of these two phases divided by the total duration of MU discharge (ascending-descending duration ratio; Figure 2A). This ratio allows the assessment of hysteresis (i.e., the amount of self-sustained discharge) with respect to force levels, with positive values indicating a leftward shift (i.e., less hysteresis) and negative values indicating a rightward shift (i.e., more hysteresis) of a single MU discharge (Hassan et al., 2021).

**Figure 2.**
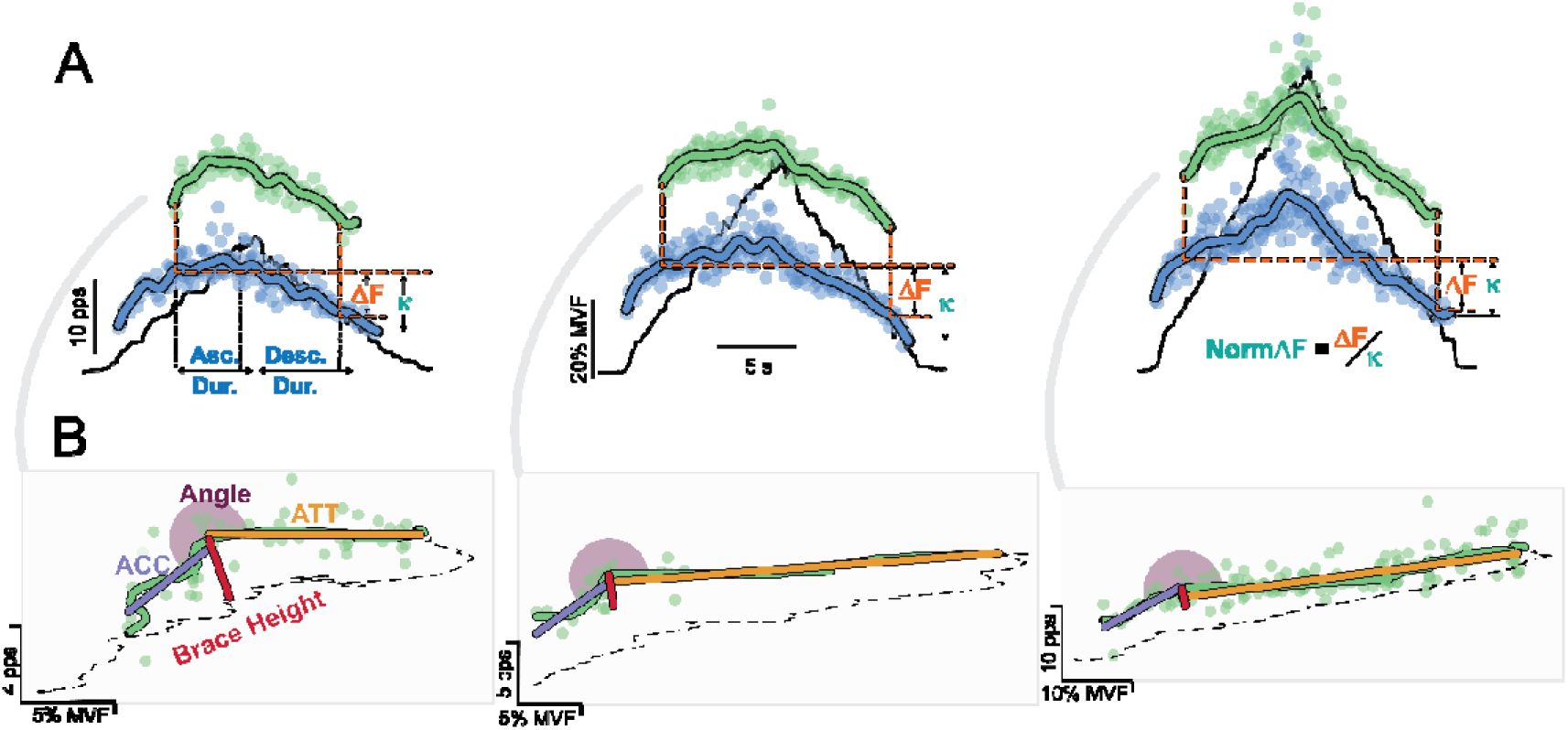
Analysis of motor unit discharge patterns. A: Paired motor unit (MU) analysis involved the calculation of onset-offset hysteresis of a higher threshold (test) MUs with respect to a lower threshold (reporter) MU (known as ΔF). For clarity, only a pair of the same MUs across three different contraction intensities (30, 50, and 70% of maximal voluntary force; MVF) is shown. Additionally, ΔF was normalised to the theoretical maximal self-sustained discharge of the test unit (the difference between the reporter unit discharge rate at test unit recruitment and reporter unit derecruitment, κ). For each MU, the ratio of the discharge duration on the ascending and descending limb of a triangular contraction was also calculated denoting the hysteresis of MU discharge with respect to peak force (negative values = more hysteresis). B: Geometric analysis of single MU discharge rate with respect to force entailed the calculation of brace height. The acceleration (ACC) and attenuation (ATT) slopes as well as the angle between the two slopes were also quantified.

##### Paired motor unit analysis

To estimate the magnitude of PICs, we calculated ΔF (Gorassini et al., 1998, 2002a), which quantifies the onset-offset hysteresis of a higher threshold (test) MU with respect to the discharge rate of a lower threshold (reporter) MU receiving common synaptic input (Figure 2A). Due to the high likelihood of many suitable reporter units for a given test unit, we calculated the average test unit ΔF for all possible reporter pairs (Hassan et al., 2021; Jenz et al., 2023). A pair of MUs was considered suitable if: 1) they exhibited a rate-rate correlation of r^2^ > 0.7 (Gorassini et al., 2004), to improve the likelihood that the pairs of MUs received common synaptic input; 2) the test unit was recruited at least 1 s after the recruitment of reporter unit, to ensure that PIC in the reporter unit were likely fully activated (Hassan et al., 2020); and 3) the reporter unit discharge rate was >0.5 pps whilst the test unit was active, to minimise the influence of MU saturation on the calculation of ΔF (Stephenson and Maluf, 2011). Additionally, we normalised ΔF values to the difference in the reporter unit discharge rate at test unit recruitment and the discharge rate at reporter unit derecruitment, as this difference is the maximal discharge rate modulation for each test-reporter unit pair; we present this as normalised ΔF, and suggest it may be an important supplement to traditional ΔF values when reporter unit discharge rates differ vastly between contractions and/or conditions. Also, note that ΔF provides an estimate of PIC magnitude but may not necessarily distinguish between the contribution of neuromodulatory drive and the amount and/or pattern of inhibitory input (Beauchamp et al., 2023; Chardon et al., 2023).

##### Geometric analysis

We used a quasi-geometric approach to estimate the deviation from linearity in MU discharge rate with respect to force output, which we previously referred to as brace height (Figure 2B; Beauchamp et al., 2023). Briefly, the smoothed discharge rate of a single MU was expressed as a function of force produced during a contraction, and the maximum orthogonal vector between this smoothed discharge and a theoretical linear increase in discharge from MU recruitment to peak discharge was quantified. The absolute magnitude of this maximum orthogonal vector is then termed brace height. To ensure that brace height quantifies a relative deviation of discharge patterns from linearity, and to remove potential confounds with changes in peak discharge, we normalised brace height values to the height of a right triangle with a hypotenuse between recruitment and peak MU discharge. MUs with a negative slope of the acceleration phase, normalised brace height greater than 200%, or with peak MU discharge rate occurring after peak force were excluded from further analysis. The estimation of brace height has been shown to be highly sensitive to changes in neuromodulatory drive in motoneuron simulations (Beauchamp et al., 2023; Chardon et al., 2023).

In addition to brace height, we calculated the slope of the initial acceleration phase of MU discharge, the subsequent attenuation phase of MU discharge, and the angle between the two phases (Beauchamp et al., 2023). The slopes of these two phases correspond to the secondary and tertiary range of MU discharge. The secondary range represents an abrupt increase in MU discharge rate following recruitment, likely due to PIC activation (Li et al., 2004), which is followed by attenuated MU discharge rate increase (tertiary range) as the PIC conductance is increased and the potassium currents are activated (Li and Bennett, 2007). The attenuation phase of the MU discharge has been shown to be particularly influenced by the *pattern* of inhibition (Powers et al., 2012; Beauchamp et al., 2023) and may thus, in combination with brace height and ΔF metrics, allow for us to uncouple the contributions of neuromodulatory and inhibitory inputs to MU discharge patterns. Conversely, acceleration phase slope and the angle between acceleration and attenuation phases have been shown to be relatively insensitive to patterns of inhibitory inputs in simulations (Beauchamp et al., 2023). However, delivering inhibitory cutaneous stimuli has previously been shown to linearise MU discharge profiles in humans (Revill and Fuglevand, 2017). Thus, rather than the pattern of inhibition, the angle between acceleration and attenuation phases is likely to be sensitive to the total inhibitory input, i.e., increased inhibitory input could lead to smaller angles.

##### Ensemble averaging

To provide a visual comparison of MU behaviour across different contraction levels, we constructed ensemble averages of all MUs identified across participants for each contraction level and muscle (Beauchamp et al., 2022). Identified MUs were separated into bins according to their recruitment threshold and the discharge patterns of MUs were smoothed with SVR as detailed above. The discharge behaviour of each MU was then normalised in time according to the average discharge duration within each ensemble cohort. This normalisation procedure involved adjusting sampling rates of the prediction vectors to equalise the length of all smoothed MU discharges within an ensemble with respect to the average length of the time vector between MU recruitment and derecruitment sampled at 2048 samples per second. Note that this approach is relevant only for the purposes of visual representation of MU behaviour and was not used when calculating any MU discharge metrics or for statistical comparisons.

#### Statistical analyses of human experiments

To assess changes in MU discharge characteristics across contraction levels, a linear mixed effects model was fit to each MU discharge metric. The model consisted of contraction level, muscle and their interaction as fixed effects and participant and trial (i.e., repeat trials) as random intercepts. Additionally, since decomposition is biased towards discriminating higher-threshold MUs in each context/contraction level (Farina et al., 2010; Francic and Holobar, 2021), and because proportions of the motor pool with moderate recruitment thresholds are likely to have been sampled across all of our contraction levels, we employed MU recruitment threshold as a covariate in our model. For example, to assess how ΔF is modulated across contraction levels in Experiment 1, our model was constructed as follows: ΔF ∼ Contraction Level*Muscle + Recruitment Threshold + [1 | Participant ID] + [1 | Trial]. For the analyses in Experiment 2, we replaced the fixed effect of ‘muscle’ with the fixed effect of ‘condition’ in our model to denote whether the contractions were performed with matched duration or matched rate of force increase. Because the 50% MVF condition was the same for both rate- and duration-matched contractions, we randomly assigned one trial at 50% to either the rate- or duration-matched condition. When assessing the change in outcome metrics across contraction levels for tracked MUs, we further considered the nesting of a particular set of tracked MUs within a given participant (e.g., peak discharge rate ∼ Contraction Level*Muscle + Recruitment Threshold + [1 | Participant ID] + [1 | Matched MU ID: Participant ID]). For ΔF data, rather than nesting of tracked MUs, we considered nesting of reporter and test units within a participant as random intercepts, as well as the recruitment threshold of test and reporter units as covariates (e.g., ΔF ∼ Contraction Level*Muscle + reporter MU threshold + test MU threshold + [1 | Participant ID] + [1 | reporter MU ID: Participant ID] + [1 | test MU ID: Participant ID]).

All analyses were performed in R (R studio, v 1.4.1106, R Foundation for Statistical Computing, Vienna, Austria). Linear mixed effect models were constructed with *lme4* package (Bates et al., 2015). The significance of the model was assessed by comparing the fit of the model with and without the predictor variables using *lmerTest* package (Kuznetsova et al., 2017). The assumptions of normality, linearity and homoscedasticity of residuals were assessed using quantile-quantile plots and histograms. For significant main effects or interactions, post hoc testing of estimated marginal means with Tukey adjustments for multiple comparisons was performed using *emmeans* package (Lenth and Lenth, 2018). Residuals of predictions were found to be non-linear for the following variables: peak discharge rate (log-transformed), discharge rate at recruitment and derecruitment (sqrt-transformed), normalised ΔF (sqrt-transformed), acceleration slope (log), and attenuation slope (log) for non-tracked analysis, and peak discharge rate (log), ascending/descending ratio (sqrt), and acceleration and attenuation slopes (both log) for tracked unit analysis in Experiment 1. For analysis of Experiment 2, the residuals of the following variables were found to be non-linearly distributed: ascending/descending ratio (sqrt), acceleration (log), and attenuation (log) for non-tracked MU data, and peak discharge rates (log), normalised ΔF (sqrt), ascending/descending ratio (sqrt), acceleration (log) and attenuation (log) for tracked MU data. In the aforementioned cases, data were transformed for the linear mixed model and then back-transformed when calculating estimated marginal means. In all cases, significance was set at an alpha level of 0.05.

### In silico experiments with realistic motoneuron pools

Previous studies using realistic motoneuron pool simulations suggested that ΔF and the slope of the ascending discharge rate modulation are sensitive indicators of the inhibitory pattern supplied to the motoneuron pool (excitation-inhibition coupling). However, these studies were limited to magnitudes of excitatory input that replicated lower-intensity contractions (Beauchamp et al., 2023; Chardon et al., 2023). Extending these metrics to higher excitation levels poses challenges because the attenuation slope is typically expressed as a function of force/torque output. Since force/torque output changes disproportionately relative to discharge rate modulation, the attenuation slope is confounded and tends to decrease at higher force levels or during rapid force production (see ‘Results’). Thus, characterising the behaviour of these metrics across excitation magnitudes at known neuromodulation and inhibitory levels is paramount to interpreting the human data presented in this work.

Moreover, earlier investigations assumed static neuromodulation regardless of excitatory conductance (Powers and Heckman, 2017; Beauchamp et al., 2023; Chardon et al., 2023). In contrast, raphe neurons dynamically adjust discharge rates based on task demands, as demonstrated by in-phase modulation during treadmill locomotion in cats (Veasey et al., 1995). Similarly, functional MRI data suggest that brainstem monoaminergic activity scales with grip strength, indicating proportional increases in monoaminergic release with excitatory input (Danielson et al., 2023). To address these gaps, we used a modelling approach with realistic motoneuron pool simulations (Powers and Heckman, 2017; Chardon et al., 2023) to examine how inhibitory patterns shape discharge rate modulation across excitation magnitudes and how neuromodulatory inputs might vary with changes in excitation.

#### Adaptations to model the human motoneuron pool

Simulations were conducted using the Python interface to the NEURON simulator with a model similar to that described previously (Powers and Heckman, 2017; available at: https://modeldb.science/239582). Unlike earlier adaptations that modelled 20 motoneurons (Powers and Heckman, 2017; Beauchamp et al., 2023; Chardon et al., 2023), this study employed an expanded pool of 35 motoneurons.

These motoneurons were assigned a range of intrinsic properties to create a gradient from low- to high-threshold motoneurons (presumably a gradient from S to FR and FF motoneurons).

Each motoneuron consisted of a soma compartment with four dendritic compartments that were parameterized with passive properties (e.g. size, capacitance, and passive conductance) as described previously by Kim and colleagues (Kim et al., 2009; Kim and Jones, 2012). Active conductances included sodium (Na) and potassium (K) channels, along with a medium afterhyperpolarization (mAHP) conductance in the soma, while dendritic compartments contained calcium (Ca) conductances to mediate the slowly activating PIC. Additionally, a hyperpolarization-activated mixed-cation conductance was distributed across all compartments.

Initial conductance densities, kinetics and steady-state activation curves were configured to recreate the input-output behaviour recorded in decerebrate cat medial gastrocnemius motoneurons (Powers and Heckman, 2017), with several adjustments to better represent human MU discharge patterns as per Chardon et al. (2023). These included: increasing the calcium removal time constant for the calcium-activated potassium conductance, resulting in more representative afterhyperpolarization durations and amplitudes; hyperpolarizing the PIC half-activation threshold to offset the modified (larger and longer) afterhyperpolarizations; and removing the voltage-dependent inactivation of PICs to capture motoneuron hysteresis behaviour more accurately.

#### The simulation protocol

To appreciate the composition of motor commands leading to the observed MU discharge patterns in human experiments, we first calculated the average cumulative spike train (CST) for TA across participants in Experiments 1 and 2. These CSTs were quantified for the triangular dorsiflexion contractions (10-second linear increase and decrease in force) to both 30% and 70% MVF. These CSTs served as reference values for the employed optimisation approach (Chardon et al., 2023).

In each simulation, a noisy Gaussian excitatory input conductance was applied to the motoneuron pool at a given neuromodulation level or inhibitory pattern. This excitatory conductance was then iteratively adjusted to align the simulated motor pool’s CST with the experimental CST. For all simulations, a baseline tonic inhibitory conductance was applied to ensure motoneurons ceased discharging in the absence of excitatory input. Additionally, excitatory conductance was scaled to ensure that the highest-threshold motoneuron (i.e. motoneuron number 35) received approximately 10% greater conductance than the lowest-threshold motoneuron.

The iterative procedure began with the experimentally observed CSTs at 30% and 70% MVF as the initial excitatory input conductance. This input was played into the motoneuron pool, and its subsequent CST was quantified. The excitatory input was then updated proportionally to the error between the simulated and experimental CSTs. This process was repeated up to 20 iterations or until convergence criteria were met, defined as achieving a mean squared error below a predefined threshold (10% peak CST) and recruiting the minimum number of motoneurons (20 for CST at 30% MVF, 35 for CST at 70% MVF).

We ran this optimization procedure independently across a range of neuromodulatory levels (multipliers of the maximum conductance of L-type calcium channels) and inhibitory patterns (scaling of inhibitory conductance relative to excitation). This included the following parameter ranges:

- Neuromodulation levels: 10 levels ranging from 0.8 to 1.7, where 1.0 corresponds to L-type calcium channel conductance in decerebrate cat motoneurons without monoaminergic drug application.
- Inhibitory conductance levels: 15 levels from -0.7 (strong reciprocal/push-pull inhibition; -0.7*excitatory conductance) to 0.7 (strong proportional inhibition), where 0 indicates tonic inhibitory conductance.

This parameter space yielded a total of 150 simulations for each CST magnitude (30% and 70% MVF).

#### The reverse engineering approach

For both contraction intensities (30% and 70% MVF), we evaluated all combinations of neuromodulation (10 levels) and inhibitory input (15 patterns) across the motor pool (∼20 motoneurons in each 30% and 35 motoneurons in each 70%). Each motoneuron’s discharge behaviour was characterised using the same metrics employed experimentally in humans: brace height, ΔF, and the ascending discharge rate modulation slope. Given that values of brace height and attenuation slope are typically quantified as a function of torque, the excitatory input for each simulation was scaled to a peak value of either 30 or 70 and used as a theoretical torque. The average value of these outcome metrics was calculated for each simulation, along with the change in these metrics across all possible simulation pairs at 30% and 70% MVF. The three-outcome metrics were chosen as they help disentangle neuromodulatory- and inhibitory-driven changes in MU discharge behaviour (Beauchamp et al., 2023).

To identify which simulated motoneuron discharge patterns most closely resembled those observed in the human experiments, we compared the in-silico values of brace height, ΔF, and attenuation slope to the experimental data. A solution space was created, including all possible 30%-70% simulation pairs that met two criteria: the outcome metrics (brace height, ΔF, and attenuation slope) had to fall within the 95% confidence intervals of the human data at both 30% and 70% MVF conditions, and the change in each metric between 30% and 70% had to align with the estimated values and 95% confidence intervals observed in the human study. To assess the accuracy of the simulated pairs that met these criteria, we calculated the Euclidean distance in three-dimensional space (brace height, ΔF, attenuation slope) between the average values observed for each simulation and the human average data. To qualitatively assess the discharge behaviour of the simulated pairs, ensembles were created that included all simulation pairs falling within the solution space, following the same procedure used to represent the human data.

## RESULTS

### Experiment 1

In the first human experiment, we identified and characterised MU discharge patterns across several contraction forces in three different human muscles (TA, VL and VM) to gain insights into the structure of motor commands. To appreciate the discharge patterns of all identified MUs across participants for each muscle, we represented MUs as ensemble averages. For each muscle, the discharge patterns of all identified MUs were smoothed and separated into ten equally spaced bins as a function of the recruitment threshold. The ensemble averages for each muscle and across all contraction levels are shown in Figure 3A, along with the mean force profile and the mean CST across all participants. The number of identified MU spike trains is presented in the Supplementary Material, both in the cases of non-tracked and tracked MUs across contraction levels (Supplemental Table 1, https://doi.org/10.6084/m9.figshare.27895032). We only present the tracked MUs below because the results were similar when assessing the discharge behaviour of all units and those that were tracked. The results of all (i.e., non-tracked) units for both experiments are presented in the Supplementary Material (https://doi.org/10.6084/m9.figshare.27895032).

**Figure 3.**
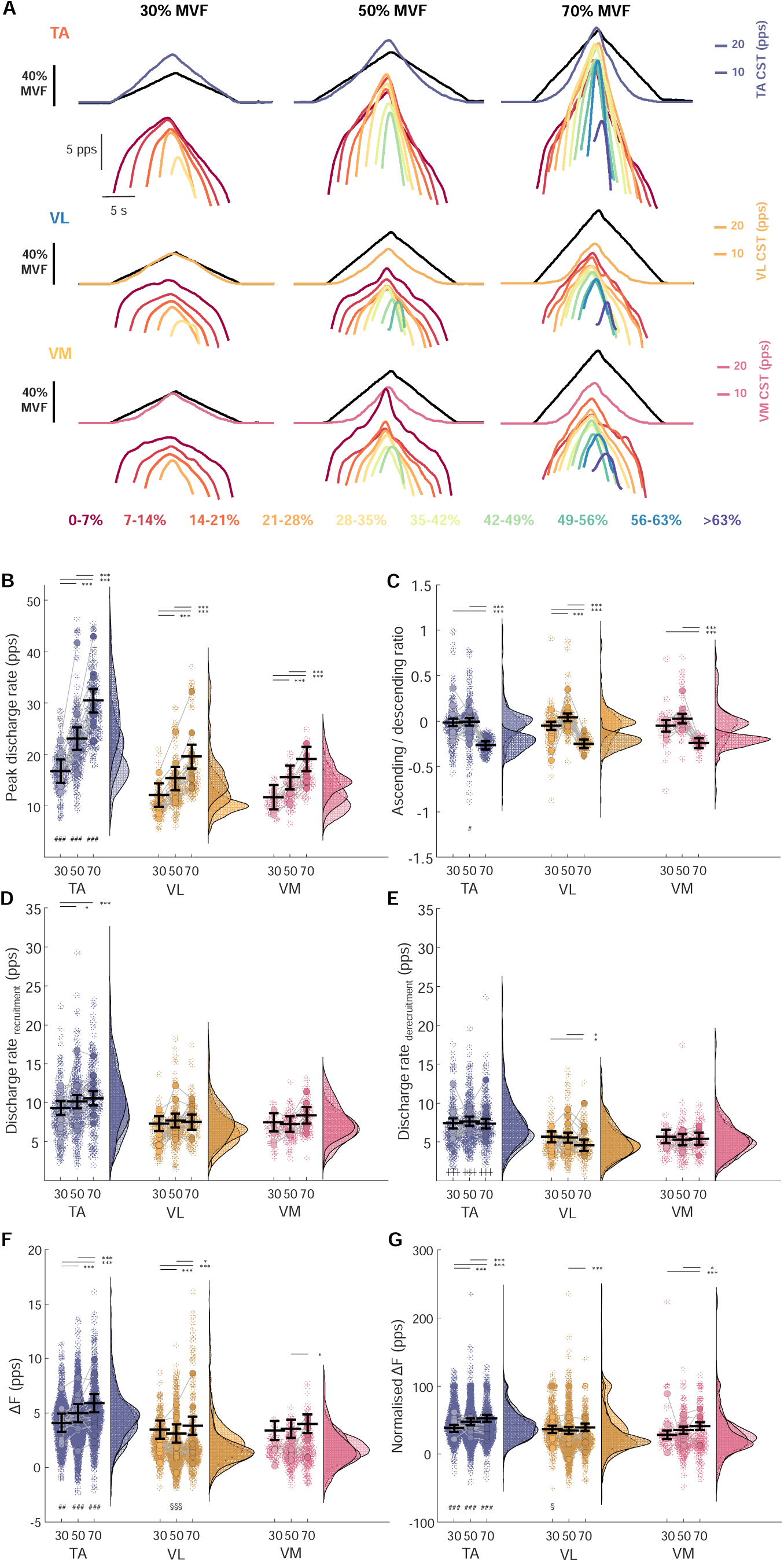
Discharge rate modulation and hysteresis as a function of contraction level and muscle. A: Ensemble averages of motor unit (MU) discharge patterns based on recruitment threshold in 7% MVF bins (colour-coded) along with the ensemble force profiles (black) and cumulative spike trains in tibialis anterior (TA), vastus lateralis (VL), and vastus medialis (VM), during triangular contractions up to 30, 50 and 70% of maximal voluntary force (MVF). B: Peak discharge rates (in pulses per second, pps) obtained as the maximal value of MU discharge patterns smoothed with support vector regression. C: the ratio of the ascending and descending phase of MU discharge rate with respect to peak force levels as an estimate of self-sustained MU discharge (positive values indicate less hysteresis). D: Discharge rate at MU recruitment. E: Discharge rate at MU derecruitment. F: Estimation of the onset-offset hysteresis (ΔF) between a pair of MUs that share a high proportion of common synaptic input. G: ΔF as in D but normalised to the to the difference in the reporter unit discharge rate at test unit recruitment and the discharge rate at reporter unit derecruitment. In B-G, the coloured circles represent individual participant averages, the opaque circles denote values of individual MUs with their kernel density distribution also plotted. The horizontal black lines denote the estimated marginal means obtained from the linear mixed statistical modelling with the error bars denoting a 95% confidence interval. ***p < 0.001, **p < 0.01, *p < 0.05 relative to other contraction levels, ^###^p < 0.001, ^##^p < 0.01, ^#^p < 0.01 relative to VL and VM at the same force level; ^§§§^p < 0.001, ^§§^p < 0.01, ^§^p < 0.05 relative to VM at the same force level; ^┼┼┼^p < 0.001 compared to VL at the same force level.

Peak discharge rates were dependent on contraction level (χ^2^(2) = 2259.4, p < 0.0001), and increased with contraction force in all muscles (p < 0.0001). We also found a significant interaction between contraction level and muscle (χ^2^(4) = 18.3, p = 0.0011), such that during contractions at the same relative force level TA peak discharge rates were greater than VL and VM (p < 0.0001; Figure 3B). The discharge rate at recruitment (Figure 3D) was also influenced by contraction level (χ^2^(2) = 19.0, p < 0.0001) and muscle (χ^2^(2) = 219.2, p < 0.0001), but there was no interaction between contraction level and muscle (χ^2^(4) = 9.5, p = 0.0507). At derecruitment (Figure 3E), discharge rate was dependent on contraction level (χ^2^(2) = 7.2, p = 0.0272), muscle (χ^2^(2) = 262.4, p < 0.0001), and recruitment threshold (χ^2^(1) = 133.8, p < 0.0001), and there was also a significant interaction between the contraction level and muscle (χ^2^(4) = 11.4, p = 0.0229).

When considering untracked MUs, there was a tendency for reduced prolongation of discharge of a single MU with respect to peak force levels with greater contraction level (see Supplemental Figure 1, https://doi.org/10.6084/m9.figshare.27895032), but this was seemingly not the case when observing the behaviour of the same units across different contraction levels. Indeed, ascending/descending ratio was influenced by contraction level (χ^2^(2) = 38.7, p < 0.0001), with a tendency for this ratio to decrease at higher contraction levels (Figure 3C) in all muscles (χ^2^(2) = 3.0, p = 0.2239), indicating a greater prolongation of MU discharge rate.

The discharge rate hysteresis (ΔF), an indicator of PIC magnitude, was dependent on contraction level, both in its original calculation (χ^2^(2) = 555.9, p < 0.0001) as well as when normalised (χ^2^(2) = 44.7, p < 0.0001) to the maximal modulation in reporter unit discharge rate possible for each test unit. In both cases, ΔF increased with contraction level (Figure 3F and 3G). There was also a significant contraction level by muscle interaction (ΔF: χ^2^(4) = 262.1, p < 0.0001; normalised ΔF: χ^2^(4) = 136.2, p < 0.0001), with ΔF typically greater in TA compared to VL and VM at the same force level (p < 0.0001).

The modulation in discharge rate in the TA with respect to force output during different contraction levels can be appreciated in Figure 4A. To appreciate the extent of non-linearity of the ascending MU discharge rate, we performed a quasi-geometric analysis of tracked units. Brace height, an indicator of non-linearity, was dependent on contraction level (χ^2^(2) = 143.2, p < 0.0001), tending to decrease with greater contraction level (Figure 4B). There was also an interaction between contraction level and muscle (χ^2^(4) = 24.9, p < 0.0001), where brace height decreased as a function of contraction level in TA (p ≤ 0.0292). In VL, brace height decreased from 30% to 50% (p < 0.0001) but then remained similar between 50 and 70% MVF (p = 1.0000), whereas in VM the only observed difference was between 30 and 70% MVF with brace height being smaller during the latter (p = 0.0046; Figure 4B). The tendency for discharge rate patterns to become more linear at higher force levels was further supported by the results of the angle between acceleration and attenuation slopes, which was dependent on contraction level (χ^2^(2) = 204.9, p < 0.0001), with a significant interaction between contraction level and muscle (χ^2^(4) = 21.2, p = 0.0003). The angle decreased as a function of contraction level in TA (p ≤ 0.0073) and VM (p ≤ 0.0425), whereas in VL there were no differences between contractions at 50 and 70% MVF (p = 0.0575; Figure 4C).

**Figure 4.**
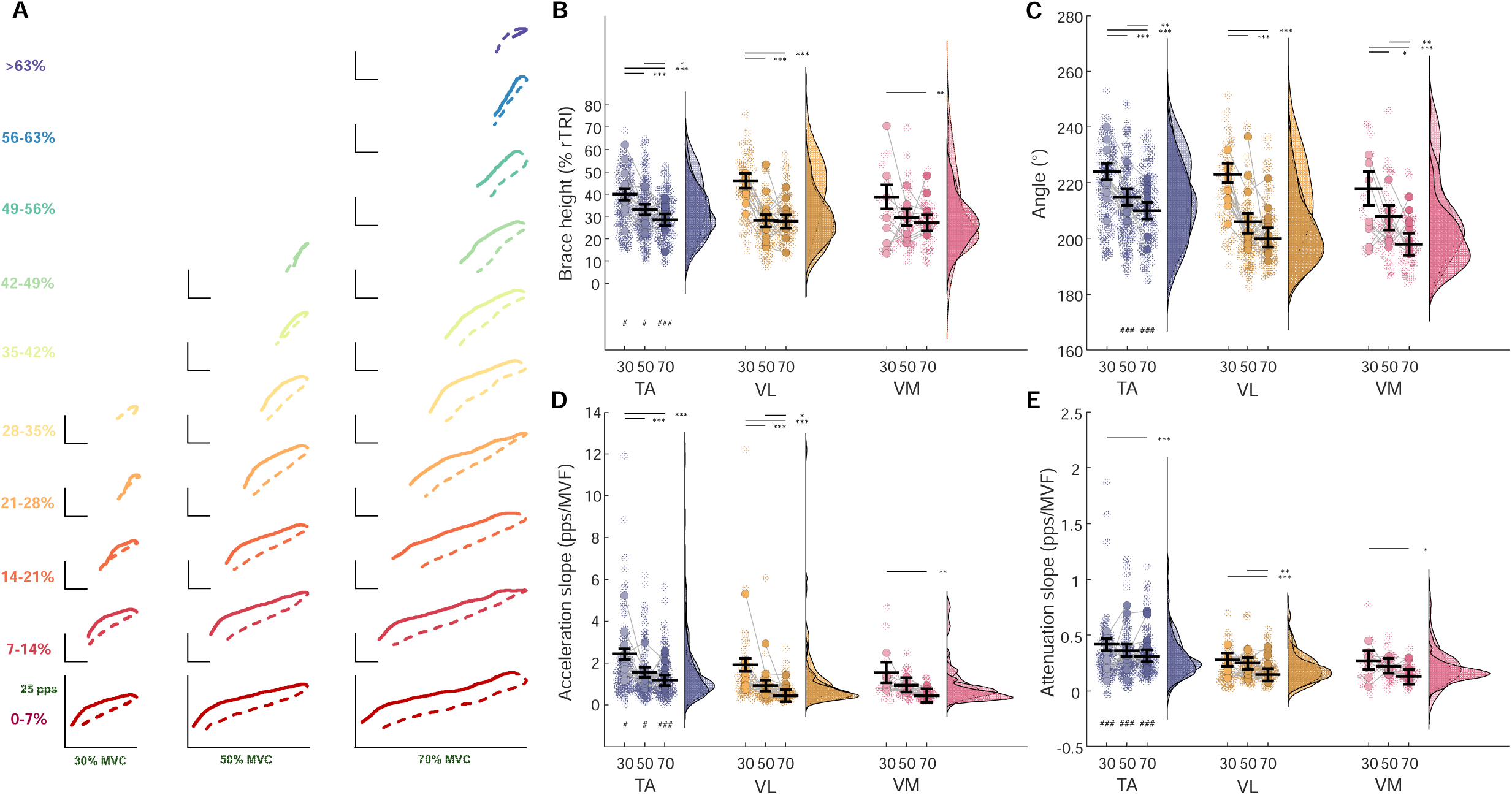
Geometric analysis of discharge rate modulation as a function of contraction level and muscle. A: Ensembles of tibialis anterior (TA) motor unit (MU) discharge (pulses per second, pps) increase (full lines) and decrease (dashed lines) as a function of a change in dorsiflexion force during 30 (left column), 50 (middle column), and 70% (right column) of MVF contractions. Each colour represents an ensemble of MU discharges stratified by recruitment thresholds in 7% MVF bins. B: Brace height expressed as a percentage of a right triangle (% rTRI) in TA, vastus lateralis (VL) and medialis (VM). C: The angle between the acceleration and attenuation slopes. D: the acceleration slope. E: the post-acceleration attenuation slope. The coloured circles represent individual participant averages, the opaque circles denote values of individual MUs with their kernel density distribution also plotted. The horizontal black lines denote the estimated marginal means obtained from the linear mixed statistical modelling with the error bars denoting a 95% confidence interval. ***p < 0.001, **p < 0.01, *p < 0.05 relative to other contraction levels, ^###^p < 0.001, ^##^p < 0.01, ^#^p < 0.01 relative to VL and VM at the same force level.

The acceleration slope, representing the secondary range of the ascending MU discharge rate, was modulated by contraction level (χ^2^(2) = 214.1, p < 0.0001), as well as contraction level and muscle interaction (χ^2^(4) = 14.4, p < 0.0001). In VL, the acceleration slope decreased with greater contraction levels (p ≤ 0.0465), whereas it also decreased between 30 and 50% for TA (p < 0.0001), but not 50 and 70% MVF (p = 0.0795). In VM, the only difference was noted between 30 and 70% (p = 0.0027; Figure 4D). The attenuation slope, denoting the tertiary range of the ascending MU discharge rate, was dependent on contraction level (χ^2^(2) = 46.5, p < 0.0001) and muscle (χ^2^(2) = 141.7, p < 0.0001). In TA and VM, the attenuation slope decreased between 30 and 70% MVF (p = 0.0001 and p = 0.0287, respectively). Conversely, in VL, the attenuation slopes were similar at 30 and 50% MVF (p = 0.9404), but then decreased between 50 and 70% (p = 0.0018; Figure 4E).

In summary, peak discharge rates increased with greater contraction force in all muscles (Figure 3B), which was accompanied by greater prolongation of MU discharge (Figure 3C) and greater discharge rate hysteresis (Figure 3F and 3G), suggesting larger PICs with greater force levels. Moreover, in all muscles examined, the ascending discharge rate tended to be more linear with greater force (brace height; Figure 4B), with lower slopes of the secondary (Figure 4D) and tertiary (Figure 4E) range of the ascending MU discharge, possibly indicating a smaller *relative* contribution of PICs to discharge rate modulation with greater excitatory synaptic input.

### Experiment 2

Due to the potential influence of contraction duration and rate of force increase on discharge rate modulation throughout the task, we performed a follow-up experiment to examine to what extent the contraction characteristics might confound the results obtained in the first experiment. Subtle differences in TA MU discharge patterns across contraction levels matched for either the contraction duration or the rate of force increase can be appreciated in the ensemble averages in Figure 5A. Note that the 50% contractions are equivalent for both duration- and rate-matched contractions.

**Figure 5.**
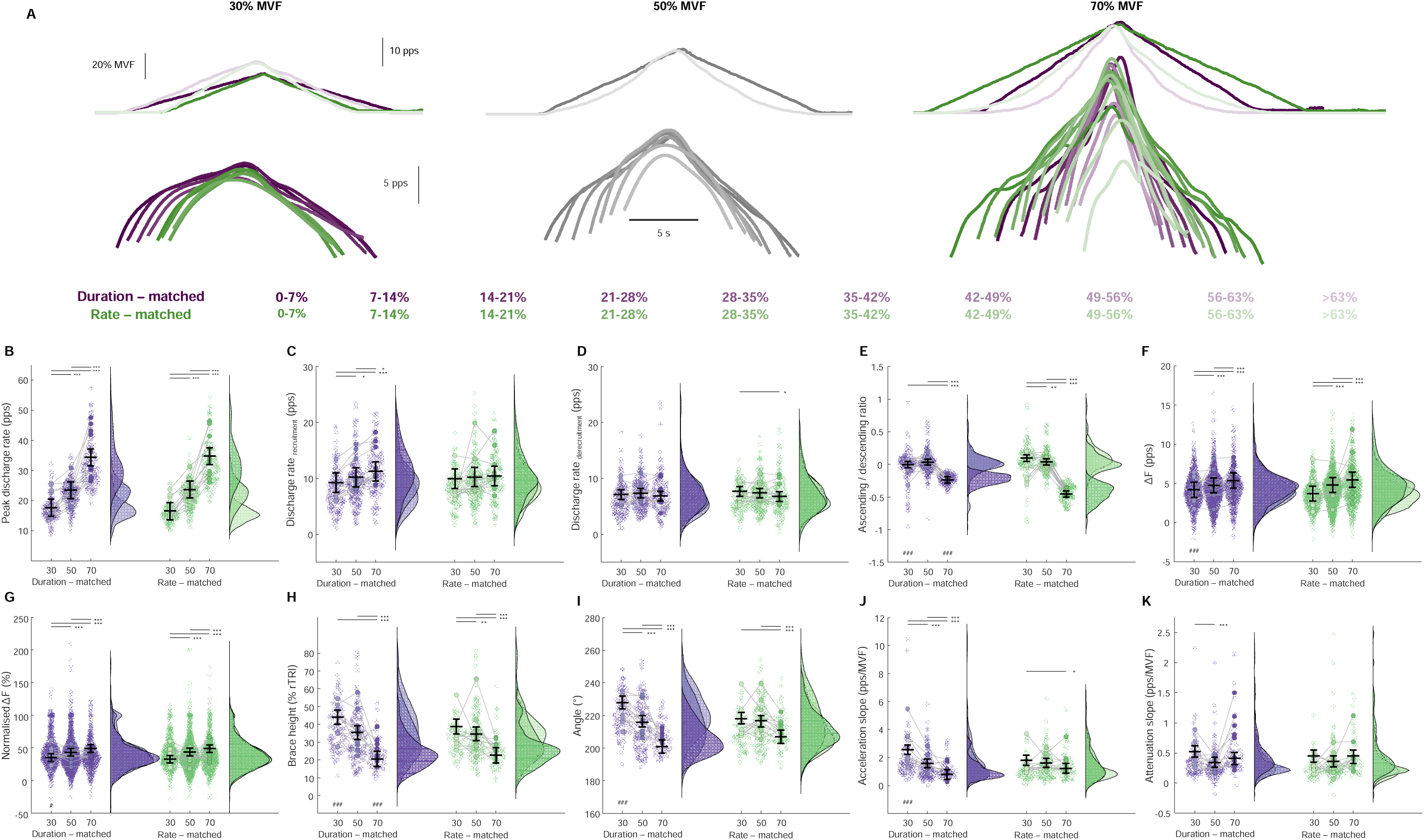
Discharge rate modulation in tracked motor units across contraction forces matched for contraction duration or rate of force increase. A: Ensemble averages of tibialis anterior motor unit (MU) discharge patterns based on recruitment threshold in 7% MVF bins (coded by the colour opacity) along with the ensemble force profiles (black) and cumulative spike trains during triangular contractions up to 30, 50 and 70% of maximal voluntary force (MVF) matched for either contraction duration or rate of force rise. B: Peak discharge rates (in pulses per second, pps) obtained as the maximal value of MU discharge patterns smoothed with support vector regression. C: Discharge rate at MU recruitment. D: Discharge rate at MU derecruitment. E: the ratio of the ascending and descending phase of MU discharge rate with respect to peak torque levels as an estimate of self-sustained MU discharge (positive values indicate less hysteresis). F: Estimation of the onset- offset hysteresis (ΔF) between a pair of MUs that share a high proportion of common synaptic input. G: ΔF as in D, but normalised to the to the difference in the reporter unit discharge rate at test unit recruitment and the discharge rate at reporter unit derecruitment. H: Brace height expressed as a percentage of the right triangle. I: The angle between the acceleration and attenuation slopes. J: the acceleration slope. K: the post-acceleration attenuation slope. In B-K, the coloured circles represent individual participant averages, the opaque circles denote values of individual MUs with their kernel density distribution also plotted. The horizontal black lines denote the estimated marginal means obtained from the linear mixed statistical modelling with the error bars denoting a 95% confidence interval. ***p < 0.001, **p < 0.01, *p < 0.05 relative to other contraction levels, ^###^p < 0.001, ^##^p < 0.01, ^#^p < 0.01 relative to the other condition at the same force level.

When considering MUs tracked across contraction levels either during duration- or rate-matched contractions, peak discharge rates were dependent on contraction level (χ^2^(2) = 2867.4, p < 0.0001), with peak discharge rates having increased with greater force level (p < 0.0001; Figure 5B), irrespective of the condition (χ^2^(1) = 2.1, p = 0.1490). The discharge rate at MU recruitment was also dependent on contraction level (χ^2^(2) = 21.1, p < 0.0001), and there was also an interaction between contraction level and condition (χ^2^(2) = 9.4, p = 0.0090), though post hoc testing did not reveal any differences between conditions at the same relative force levels (Figure 5C). Discharge rate at MU derecruitment was also dependent on contraction level (χ^2^(2) = 9.7, p = 0.0079), but was not influenced by condition (χ^2^(1) = 0.9, p = 0.3310; Figure 5D).

The ascending/descending duration ratio was influenced by contraction level (χ^2^(2) = 28.5, p < 0.0001), condition (χ^2^(1) = 10.2, p = 0.0014), as well as contraction level and muscle interaction (χ^2^(2) = 7.4, p = 0.0065). For rate-matched contractions, ascending/descending duration decreased with increased contraction level (p ≤ 0.0084), whereas for duration-matched there was an overall reduction in the ascending/descending duration with contraction level but no difference between 30 and 50% MVF (p = 0.1198; Figure 5E), suggesting that the rate of increase/decrease in force plays an important role in the quantification of MU discharge prolongation.

Discharge rate hysteresis was dependent on contraction level (ΔF: χ^2^(2) = 554.9, p < 0.0001, normalised ΔF: χ^2^(1) = 436.2, p < 0.0001), and there was an interaction between contraction level and condition (ΔF: χ^2^(2) = 64.8, p < 0.0001, normalised ΔF: χ^2^(2) = 8.3, p = 0.0158). Specifically, discharge rate hysteresis increased with greater contraction level (p < 0.0001), with greater values for the duration-matched condition at 30% MVF where the total contraction duration was longer (ΔF: p < 0.0001, Figure 5F; normalised ΔF: p = 0.0181, Figure 5G).

Brace height, an index of non-linearity of the ascending MU discharge rate, was influenced by contraction level (χ^2^(2) = 216.4, p < 0.0001), but not condition (χ^2^(2) = 3.5, p = 0.0629). There was an interaction between contraction level and muscle (χ^2^(2) = 11.2, p = 0.0038). In duration-matched contractions, brace height decreased with greater contraction level (p < 0.0001), whereas, although it was reduced at 70%, it was similar between 30 and 50% MVF for rate-matched contractions (p = 0.0503; Figure 5H). The angle between acceleration and attenuation slopes was dependent on contraction level (χ^2^(2) = 183.9, p < 0.0001), and there was a significant interaction between contraction level and condition (χ^2^(2) = 41.8, p < 0.0001). In duration-matched contractions, the angle decreased as a function of contraction level (p < 0.0001), whereas during rate-matched contractions, the angle was similar at 30 and 50% MVF (p = 0.9310). At 30% MVF, the angle was greater for longer contractions (p < 0.0001; Figure 5I).

Acceleration slope was influenced by contraction level (χ^2^(2) = 106.0, p < 0.0001) and recruitment threshold (χ^2^(1) = 84.2, p < 0.0001), but not condition (χ^2^(2) = 2.6, p = 0.1077). A significant interaction between contraction level and condition was also observed (χ^2^(2) = 41.7, p < 0.0001). In duration-matched contractions, acceleration slope decreased with greater contraction level, whereas they were only smaller during 70% compared to 30% MVF for the rate-matched contraction (p = 0.0132; Figure 5H). The acceleration slope was also greater during duration-matched compared to rate-matched at 30% MVF (p < 0.0001; Figure 5J), indicating that PIC activation played a larger role in the initial discharge rate increases during slow rates of force increase.

Attenuation slope was dependent on contraction level (χ^2^(2) = 25.2, p < 0.0001) and recruitment threshold (χ^2^(1) = 80.7, p < 0.0001), but not condition (χ^2^(1) = 0.1, p = 0.8185; Figure 5K) suggesting that the rate of increase in force plays a significant role in the estimation of these measures and their ability to draw inferences about mechanisms underlying the modulation of MU discharge patterns across intensities.

Collectively, the greater MU discharge rate prolongation (Figure 5E), discharge rate hysteresis (Figure 5F and 5G), and more linear ascending MU discharge rate modulation (Figure 5H) with greater contraction force were evident regardless of whether the contractions were matched for duration or rate of force rise. In pairwise comparisons, longer contractions tended to exhibit greater discharge rate hysteresis at the same force level, likely due to spike-frequency adaptation (Revill and Fuglevand, 2011; Vandenberk and Kalmar, 2014), and greater non-linearity of the ascending discharge rate modulation, likely related to the slow activation of PICs (Lee and Heckman, 1998b). Finally, whilst the slopes of the secondary (Figure 4D) and tertiary (Figure 4E) range of the ascending MU discharge decreased with greater contraction force for the duration-matched contractions, they remained largely unaffected by contraction force for rate-matched contraction.

### In silico experiments with realistic motoneuron pools

For each excitatory conductance, we ran 150 simulations across neuromodulation levels and inhibitory patterns. Each lower excitatory conductance level (30% MVF) simulation included ∼20 motoneurons, yielding 3,000 simulated motoneurons, while the higher excitatory conductance level (70% MVF) simulations included ∼35 motoneurons, yielding 4,500 motoneurons. The general optimisation procedure is shown in Figure 6A, and the average values for brace height, ΔF, and attenuation slope are presented in Figure 6B. These values represent the average across all motoneurons in a given simulation and illustrate the relationship between each outcome metric, neuromodulation level (on the x-axis), and inhibitory pattern (colour- coded).

**Figure 6.**
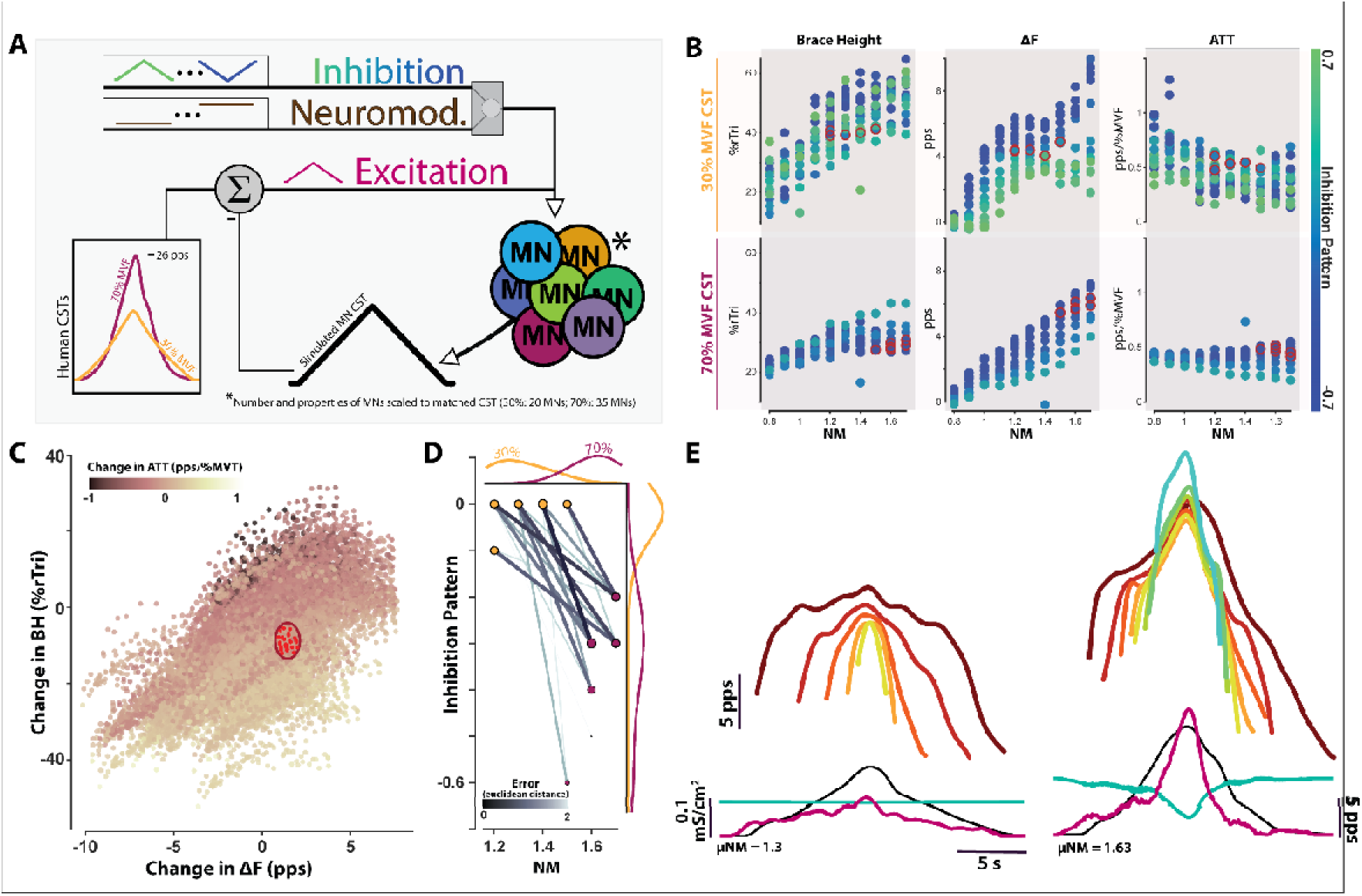
Changes observed in human motor units from lower to higher contraction forces are reproduced in silico by a more reciprocal pattern of excitation-inhibition coupling and greater neuromodulatory levels. A: The employed optimization framework iteratively adjusted a noisy excitatory conductance to align the cumulative spike trains (CSTs) of simulated motoneurons with those from experimental data for 30% and 70% maximal voluntary force (MVF) contractions. Convergence required minimal motoneuron recruitment (20 for 30%, 35 for 70%) and a mean squared error of less than 10% of the peak CST value. Simulations were performed across 10 neuromodulation (NM) levels [0.8 – 1.6] and 15 inhibitory patterns [-0.7 – 0.7]. Values of NM indicate a multiplier applied to the Ca^2+^ conductance mediating persistent inward currents, and inhibition values indicate the relation between excitatory and inhibitory conductance provided to the motoneuron. B: Average values of the outcome metrics brace height (BH), discharge rate hysteresis (ΔF), and attenuation slope (ATT). Conditions circled in red indicate those that exhibit values falling within the 95% confidence intervals of the human data for each intensity condition. C: The change in outcome metrics between all possible low and high excitatory conductance simulation pairs with the red ellipse indicating the 95% confidence interval for the changes observed in the human data set. Red dots indicate the 19 pairs that match the human data. D: The NM level and inhibitory patterns for the low (yellow) and high excitatory conductance (pink) simulation pairs that matched the changes observed in the human dataset. Probability density functions are displayed along the x- and y-axis for each intensity. Colour saturation and width of the lines connecting pairs indicate the error (Euclidean distance) between the changes observed in the simulated and human data. E: The average discharge behaviour for the low and high-intensity simulations that matched the human data, on the left and right respectively. Ensemble averages were generated in cohorts based on recruitment threshold in 7% MVF bins (colour-coded), with the average CST (black trace), average inhibitory conductance (teal trace), and average excitatory conductance (pink trace) shown. pps: pulse-per-second; %rTri: percentage of a right triangle.

For the low excitatory conductance level simulations, all combinations achieved convergence. However, for the higher excitatory conductance level simulations, convergence was not possible for inhibition patterns above 0.2. While outcome metrics generally followed trends reported previously for 30% MVF contractions (Beauchamp et al., 2023; Chardon et al., 2023), trends for the 70% MVF simulations appear dampened, partly explained by the lack of convergence with proportional (+) inhibition patterns.

Across all neuromodulation levels and inhibitory patterns, 6 combinations in the lower excitatory conductance simulations yielded outcome metrics that fell within the 95% confidence intervals for the values observed in the human experiments, with an average neuromodulation level of 1.3 and an average inhibition pattern of -0.07 (neuromodulation level range: [1.1, 1.5]; inhibition pattern range: [-0.30, 0.0]). Similarly, 8 conditions in the higher excitatory conductance simulations exhibited outcome metrics that aligned with those observed in the 70% MVF contractions, with an average NM level of 1.6 and inhibition pattern of -0.45 (neuromodulation level range: [1.5, 1.7]; inhibition pattern range: [-0.70, -0.20]).

The change in outcome metrics between all possible low- and high excitatory conductance simulation pairs is shown in Figure 6C. The red ellipse highlights the 95% confidence interval of changes observed in the human data from 30% MVF to 70% MVF (brace height: [-14.2, -6.5], ΔF: [0.910, 2.06], attenuation slope: [-0.20, - 0.02]), marking the 19 possible simulation pairs that fell within this range. The change in neuromodulation levels and inhibitory patterns between these 19 pairs are depicted in Figure 6D, with the line saturation representing the Euclidean distance in three-dimensional space (brace height, ΔF, attenuation slope) between the change in outcome metrics observed in the human data and those from the simulation pair. Across all matched pairs, the average distance was 1.3 ± 0.8, with the lowest distance of 0.14 observed between a 30% simulation at 1.4 neuromodulation and 0.0 inhibition, and a 70% simulation at 1.6 neuromodulation and -0.3 inhibition.

For all matched pairs, the directionality of neuromodulation and inhibitory pattern changes were consistent, with all 19 pairs exhibiting a more reciprocal/push-pull (-) pattern of excitation-inhibition coupling and a greater level of neuromodulation. On average, to reproduce the discharge relationships observed in the human data, in silico motoneurons needed to be supplied an excitation-inhibition coupling pattern that was more negative by -0.3 ± 0.1 and an increased neuromodulation level of 0.33

## DISCUSSION

Leveraging the combined approach of multichannel EMG decomposition with a high yield of MU discharge patterns and computer models of realistic motoneuron pools, we show that neuromodulation and patterns of inhibition are uniquely shaped to support greater contraction force across a large proportion of the motor pool’s recruitment range. Specifically, our results suggest that while the absolute magnitude of PICs increases with greater excitatory input, their relative contribution to discharge rate acceleration immediately after recruitment (i.e., brace height, PIC acceleration) decreases as a function of the magnitude and/or the rate of increase in excitatory input to human motoneurons. Concomitantly, our data indicate that modulation of inhibition facilitates force production, becoming more reciprocal relative to excitatory synaptic inputs as the demand for force increases.

### Onset-offset hysteresis increases as a function of contraction force

In agreement with our hypothesis, the observation of a greater onset-offset hysteresis (i.e., ΔF) and sustained discharge at higher contraction levels suggests that the contribution of PICs to the prolongation of motoneuron discharge increases with force output. These results are consistent with prior reports in the TA and gastrocnemius medialis muscles (Orssatto et al., 2021; Goodlich et al., 2023), but contrast those reports that showed no differences in discharge rate hysteresis with increased contraction levels in TA and soleus (Afsharipour et al., 2020; Kim et al., 2020). Though the reason for these conflicting findings is difficult to discern, methodological factors are likely contributors (e.g. the choice of smoothing function (Beauchamp et al., 2022), pairwise vs. unit-wise approach (Hassan et al., 2020), a composite reporter unit vs. individual reporter-test unit pairs (Afsharipour et al., 2020), as is the range of contraction forces, and thus recruitment thresholds, which, when small (usually 10-30% MVF; Afsharipour et al., 2020; Kim et al., 2020), make it difficult to detect small differences in ΔF.

The greater onset-offset hysteresis at higher contraction force is unlikely due to the greater discharge rates as the observations were preserved when ΔF was normalised to the maximal modulation in reporter unit discharge rate possible for each test unit (see Figure 2B). Furthermore, in agreement with our second hypothesis, the relative increase in onset-offset hysteresis with greater contraction level was maintained when contractions were matched for either total duration or the rate of force increase (Experiment 2), suggesting that the effect of contraction force on discharge rate prolongation is unimpeded by intrinsic motoneuron properties such as spike frequency adaptation and spike threshold accommodation (Bradley and Somjen, 1961; Miles et al., 2005b; Vandenberk and Kalmar, 2014). Finally, though the relative contribution of monoaminergic inputs likely varies between motor pools (Hounsgaard et al., 1988; Wilson et al., 2015; Avrillon et al., 2024), the inconsistencies in the effects of contraction force on estimates of PICs reported in the literature are unlikely due to differences in discharge rate hysteresis in different motor pools, as discrepant results have been found even when examining the same muscle (Kim et al., 2020; Orssatto et al., 2021). In support of this supposition, we observed an increasing trend of discharge rate hysteresis with greater contraction force regardless of muscle, suggesting that self-sustained discharge is contraction force dependent.

### Amplification versus prolongation of motoneuron discharge

The input-output relationship of motoneurons governs the discharge characteristics of MUs and is dependent on the intrinsic electrical properties of motoneurons, as well as the pattern and relative contribution of synaptic excitation, inhibition, and neuromodulation (Johnson et al., 2017b). Indeed, the observed increase in onset-offset hysteresis with greater contraction force suggests an increase in PIC magnitudes with a greater contraction level. Considering the dependence of PIC magnitude on monoaminergic release (e.g., 5HT; Lee and Heckman, 1998a, 2000), the observed behaviour is consistent with the notion of monoamine-mediated (i.e., 5HT and noradrenaline) gain control mechanism of spinal motoneuron discharge to support greater force production (Wei et al., 2014). Further considerations are required, however. First, the monoaminergic release tends to be slow (White et al., 1991; Raymond et al., 2001; Hentall et al., 2006) and thus the comparison with pharmacological studies that substantially alter the cellular availability of monoamines is challenging. Second, discharge rate hysteresis (ΔF) largely assesses a particular property of PICs, that is – prolongation of MU discharge (mediated predominantly by persistent Ca currents; Lee and Heckman, 1998b; Bennett et al., 2001; Li and Bennett, 2003; ElBasiouny et al., 2006; Harvey et al., 2006; Moritz et al., 2007). However, ΔF is less likely to be sensitive to the remaining hallmarks of PIC action on MU discharge, namely, amplification (Lee and Heckman, 2001; Harvey et al., 2006; Kuo et al., 2006), and rate attenuation (Powers and Heckman, 2017). Finally, neuromodulatory inputs are highly diffuse and thus a local and specific inhibitory system is required to fine-tune PIC activation (Johnson and Heckman, 2014).

Despite the observed increases in ΔF with increased contraction force, the MU discharge rate non-linearity (brace height; Figure 1C) decreased. In simulations with a single level of excitatory conductance, brace height is tightly predictive of changes in neuromodulation and is, unlike ΔF, seemingly uninfluenced by the pattern of inhibitory inputs (Beauchamp et al., 2023; Chardon et al., 2023). We extend these findings in our simulations by providing evidence that the MU discharge rate non- linearity appears to be linked to neuromodulatory drive at two different levels of excitatory conductance (Figure 6B). The deviation in linearity that brace height represents is likely the result of monoaminergic-mediated PIC facilitation and thus amplification of motoneuron discharge (Powers and Heckman, 2017), which is due to their rapid activation and inactivation properties allowing the generation of inward currents (Lee and Heckman, 2001; Harvey et al., 2006; Kuo et al., 2006). The greater linearity of discharge patterns with greater contraction force indicates a *relatively* smaller contribution of PICs to the ascending motoneuron discharge when excitatory synaptic input is disproportionally larger. However, the actions of PICs are time-dependent, with sufficient time required for full activation. This is evident when comparing triangular contractions of different durations, but similar levels of force output, whereby longer contraction durations resulted in less linear discharge rate modulation (i.e., greater brace height) compared to shorter contractions (Experiment 2, Figure 4H). Nevertheless, the trend of discharge patterns becoming more linear with greater contraction force was maintained even when accounting for the rate of force increase, suggesting that both the magnitude and the rate of increase in excitatory synaptic input influence the relative contribution of neuromodulatory inputs to the linearity of motoneuron discharge.

### Uncoupling the relative contribution of neuromodulation and patterns of inhibition to motoneuron discharge

In addition to neuromodulatory inputs, the extent and pattern of inhibition relative to excitation is an important consideration when interpreting our data. The large number and density of inhibitory synapses near voltage-sensitive channels responsible for PICs suggest a potent effect of inhibitory inputs on PIC-mediated gain control (Johnson and Heckman, 2014). The inhibitory inputs are thought to either covary with (proportional inhibition; Berg et al., 2007), or reduce in contrast to (reciprocal/push-pull inhibition; Johnson et al., 2012) excitatory synaptic input. In our simulations with excitatory conductances that reproduced the CSTs of 30% and 70% contractions, we demonstrated that the relative changes in onset-offset hysteresis, ascending discharge rate non-linearity (brace height), and attenuation slopes could only be replicated by 19 possible combinations of neuromodulation (magnitude of L-type calcium channel conductance) and patterns of inhibitory conductance. In all cases, inhibitory patterns decreased from a tonic baseline with greater excitatory input (i.e., reciprocal inhibition-excitation coupling) and neuromodulatory drive increased. Though the source of these inhibitory inputs is elusive and requires further investigation, reciprocal Ia inhibition (Hyngstrom et al., 2007) and recurrent (Renshaw cell) inhibition (Bui et al., 2005) are thought to be potent mediators of PIC contribution to motoneuron discharge. Nevertheless, the combination of human experimental and in silico data provides strong evidence that the greater PIC persistence at greater contraction forces (ΔF) was likely induced by not only a greater neuromodulatory drive, but also inhibitory patterns that became more reciprocal in nature with increasing excitatory input.

### Conclusions

Modifications in MU behaviour across contraction forces appear to be mediated by alterations in motor commands (i.e., neuromodulation and the pattern of excitatory and/or inhibitory synaptic inputs). As contraction force is increased, we found an increase in self-sustained discharge, likely facilitated by calcium PICs, and more linear MU discharge patterns. A *multi*-metric analysis approach, combined with computer simulations of realistic motoneuron pools, suggests that these observations could be mediated by more reciprocal patterns of inhibition relative to excitatory inputs and larger PICs. These data suggest that modulation of contraction force is orchestrated by a uniquely shaped combination of excitatory, neuromodulatory, and inhibitory synaptic inputs that influence the intrinsic properties of motoneurons underlying MU discharge patterns.

## Acknowledgements

The authors thank Prof CJ Heckman for useful discussions, providing comments on a draft of the manuscript, and for sharing the motoneuron model. The authors are also grateful to Miss Tamara Valenčič, Mr Chris Connelly, and Mr Haydn Thomason for assistance with data collection.

J.Š. was supported by Versus Arthritis Foundation Fellowship (reference: 22569). G.E.P.P. was supported by the Natural Sciences and Engineering Research Council of Canada (Discovery Grant RGPIN-2023-05862, and Discovery Launch Supplement DGECR-2023-00279).

